# microRNA-triggered transposon small RNAs mediate genome dosage response

**DOI:** 10.1101/203612

**Authors:** Filipe Borges, Jean-Sébastien Parent, Frédéric van Ex, Philip Wolff, German Martínez, Claudia Köhler, Robert A. Martienssen

## Abstract

Chromosome dosage plays a significant role in reproductive isolation and speciation in both plants and animals, but underlying mechanisms are largely obscure^1^. Transposable elements can promote hybridity through maternal small RNA^2^, and have been postulated to regulate dosage response via neighboring imprinted genes^3,4^. Here, we show that a highly conserved microRNA in plants, miR845, targets the tRNA^Met^ primer-binding site (PBS) of LTR-retrotransposons in *Arabidopsis* pollen, and triggers the accumulation of 21 to 22-nucleotide small RNA in a dose dependent fashion via RNA polymerase IV. We show that these epigenetically activated small-interfering RNAs (easiRNAs) mediate hybridization barriers between diploid seed parents and tetraploid pollen parents (“the triploid block”), and that natural variation for miR845 may account for “endosperm balance” allowing formation of triploid seeds. Targeting the PBS with small RNA is a common mechanism for transposon control in mammals and plants, and provides a uniquely sensitive means to monitor chromosome dosage and imprinting in the developing seed.

---

Epigenetic silencing of transposable elements (TEs) is regulated by cytosine methylation (mC), which is established by RNA-directed DNA methylation (RdDM) and occurs in three sequence contexts in plants (mCG, mCHG and mCHH, where H is A, C or T)^5^. TE methylation can be maintained independent of RdDM, but undergoes reprogramming in the germline via small RNA and histone modification, in animals and to some degree in plants^6^. Flowering plants reproduce sexually through double fertilization, resulting from the fusion of two sperm cells (in pollen grains) with the egg cell and central cell (in the embryo sac), to produce a diploid embryo and a triploid endosperm, respectively^7^. In *Arabidopsis* pollen^8–10^ and endosperm^11,12^ active removal of DNA methylation from TEs flanking imprinted genes is essential for fertility and for seed viability^10,13^. Targeted demethylation occurs specifically in vegetative and central cell nuclei and has been associated with the accumulation and mobilization of easiRNA into neighboring germ cells^8–10,14^, but the biological significance of these small RNAs has remained unclear. Here we show that easiRNAs play a critical role in epigenetic reprogramming before and after fertilization in interploid hybrids.

easiRNA from TE transcripts are triggered by microRNAs (miRNAs), most notably miR845a (21-nt) and miR845b (22-nt) that target multiple transposons in Arabidopsis pollen^15^ (Fig. 1a), where TEs are also expressed^9^. miR845a and miR845b target specific *Gypsy* and *Copia* retrotransposons at the 18nt primer-binding site (PBS) (Fig. 1b and Supplementary Table 1), where tRNAs initiate reverse transcription, and their targets generate easiRNA specifically in pollen (Fig. 1c). *MIR845* belongs to a highly conserved family of miRNAs in plants with a complex history of retention and loss in Brassicaceae^16^, and is not present in the perennial *Arabis alpina* where *Gypsy* transposons otherwise targeted by miR845 have contributed to a massive genome expansion^17^. *MIR845* orthologs have been found in drought-stressed rice leaves^18^ and diploid strawberry^19^, and may have been derived from tRNA^iMet^ through a single inversion event^19^. However, as target complementarity extends from the PBS towards the 5’ LTR (Fig. 1b), it appears that *MIR845* in Arabidopsis has derived from truncation and inversion of 5’ LTRs (plus PBS). Intriguingly, in mammalian cells, abundant 18 to 22-nt 3’CCA tRNA fragments also match endogenous retroviruses at the PBS, and strongly suppress retrotransposition, suggesting an ancient mechanism for transposon control^20^.

**Figure 1.**
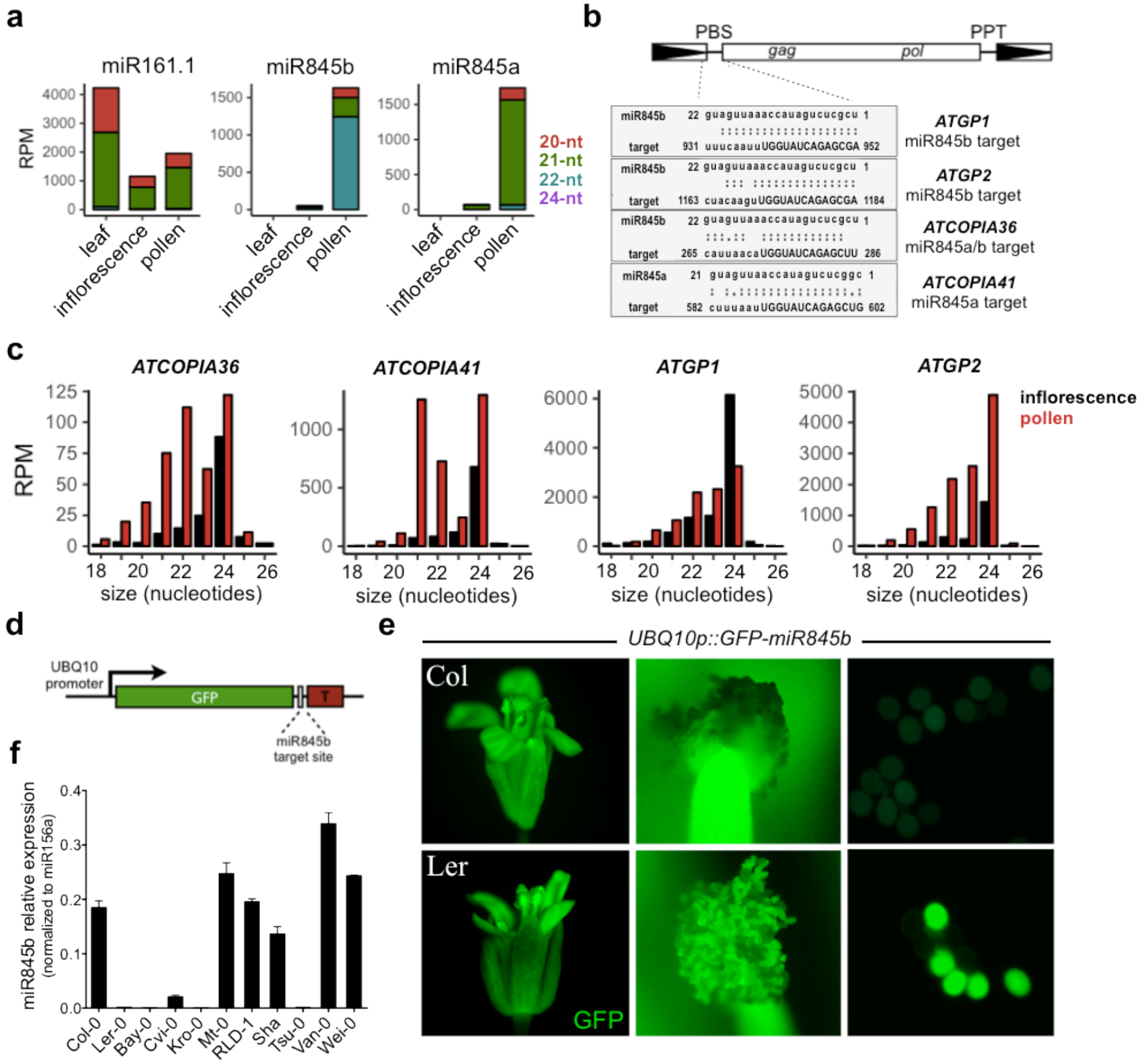
miR845 family is expressed in pollen and targets retrotransposons. **a**, miR845a and miR845b are preferentially expressed in mature pollen, consistent with complete absence in leaves and low levels in inflorescence tissue. **b,** miR845 targets the PBS (uppercase) of retrotransposons where tRNAs bind to initiate reverse transcription. Predicted targets include *Gypsy* and *Copia* elements. **c**, miR845 targets accumulate high levels of secondary 21- and 22-nt easiRNA in pollen. **d**, GFP sensor construct includes 3’UTR with a miR845b target site and driven by the *UBIQUITIN10* (*UBQ10*) promoter. **e,** Strong GFP fluorescence was detected in floral organs, but not in wild type Col-0 pollen. The same reporter is not silenced in Ler-0 pollen, where miR845b is not expressed. **f**, The *MIR845* haplotype in Ler-0 is also found inother Arabidopsis accessions such as Bay-0, Kro-0 and Tsu-0 that produce low levels of miR845b, while Col-like accessions express high levels of miR845b.

We used a GFP sensor incorporating a 3’UTR miR845b target site and driven by the ubiquitously expressed *UBIQUITIN10* (*UBQ10*) promoter (Fig. 1d) to demonstrate strong reduction of GFP expression in pollen (Fig. 1e) and sperm cells (Supplementary Fig. 1a) of the Arabidopsis accession Columbia (Col-0). Strikingly, the same GFP sensor was not silenced in pollen from Landsberg (Ler-0), suggesting that miR845b was missing from this accession (Fig. 1e). In Ler-0, *MIR845a* has been deleted completely while *MIR845b* has a single nucleotide polymorphism (SNP) in the complementary miRNA* sequence that creates an extra bulge in the duplex that impairs miRNA processing (Supplementary Fig. 2a-f). Importantly, the Ler-0 *MIR845* haplotype is conserved in many Arabidopsis accessions such as Kro-0, Bay-0 and Tsu-0, and we found that the levels of miR845b are depleted in these ecotypes (Fig. 1f). To address the potential effect of miR845 on TE silencing, we compared Col-0 and Ler-0 pollen transcriptomes and found that TE transcripts are overall more abundant in Ler-0 pollen (Supplementary Fig. 3a), including miR845 targets such as *ATGP2* and *ATCOPIA36* that have escaped silencing in this ecotype (Supplementary Fig. 3c). Further, miR845-targeted TEs such as *ATGP2* and *ATCOPIA41* accumulate easiRNA in Col-0 pollen, while other TE families such as *ATCOPIA63* are expressed and produce easiRNA specifically in Ler-0 (Supplementary Fig. 3b).

easiRNAs have been associated with RdDM activity^15,21^, and in pollen, these pathways are activated in the vegetative cell nucleus (VN) but not in sperm cells (SC)^8^, where 21 and 22-nt easiRNAs accumulate^9^. Therefore we performed bisulfite sequencing of FACS-sorted Col-0 and Ler-0 pollen nuclei, and found that levels of mCHH in the Ler-0 VN were substantially lower than in Col-0 VN (Fig. 2a), resembling seed methylomes in this respect^22^. This difference is particularly striking in pericentromeric heterochromatin, where the chromomethylases CMT2 and CMT3 maintain mCHH and mCHG, respectively (Fig. 2a). In contrast, the levels of mCHH in Col-0 and Ler-0 SC nuclei were found to be identical and very low (Fig. 2a)^8,10,23^. In the VN, we found approximately 6000 differentially methylated regions (DMRs) for mCHH between Col-0 and Ler, which overlapped primarily with TEs (including predicted miR845 targets) (Fig. 2b and Supplementary Table 2). The majority of DMRs were hypermethylated in Col-0 VN (Fig. 2b), but not in the VN methylomes of the RdDM mutant *nrpd1a* (largest subunit of RNA Polymerase IV) or *cmt2*, indicating the that loss of mCHH in Ler-0 VN occurs at particular loci targeted by CMT2 and Pol IV-dependent siRNAs (Fig. 2c-e). Taken together, these observations suggest that natural variation in miR845 and easiRNA biogenesis is associated with deregulation of TE silencing in pollen, which might have contributed to epigenetic variation in Arabidopsis populations^24^.

**Figure 2.**
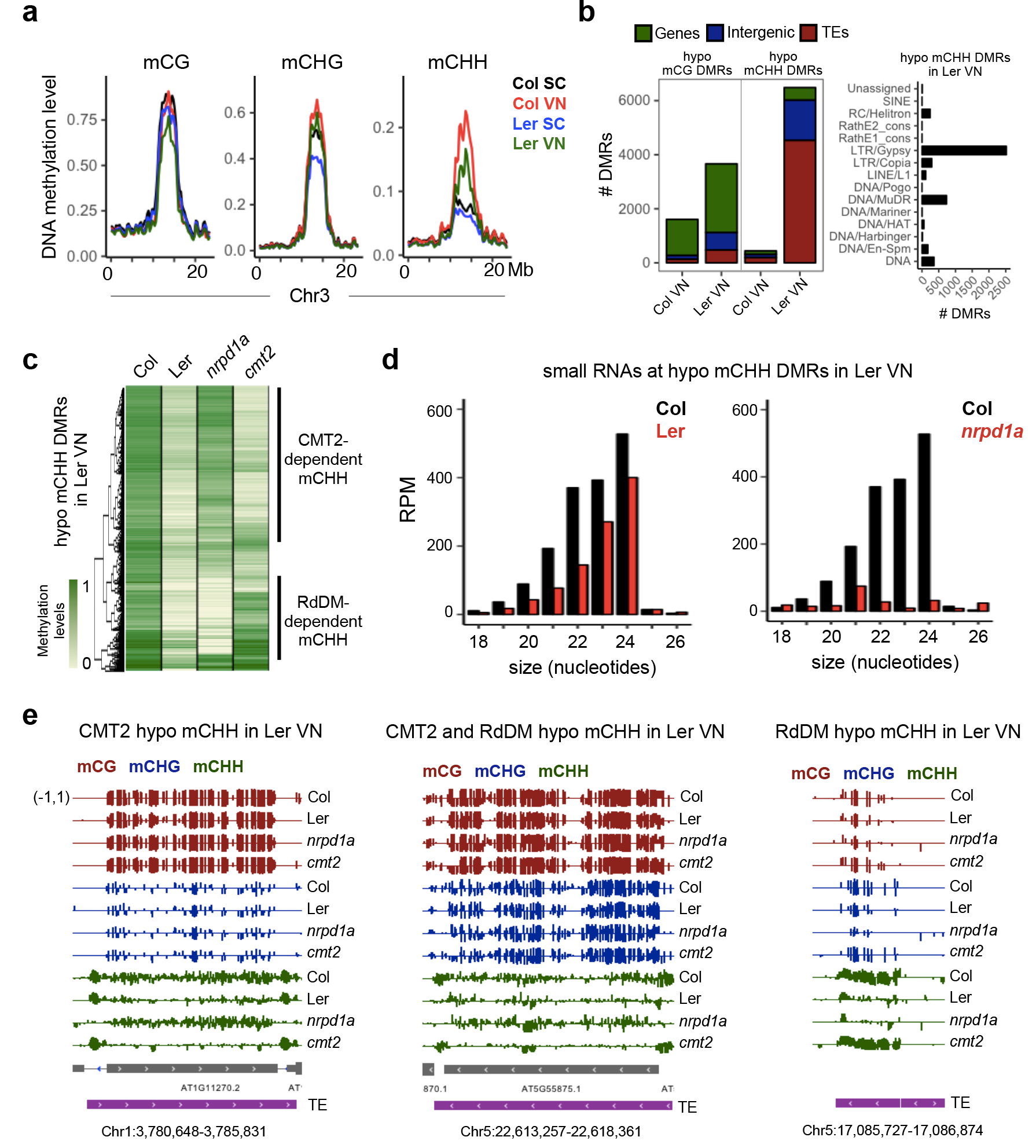
Natural variation in DNA methylation levels in Col-0 and Ler-0 pollen nuclei. **a**, Bisulfite sequencing of FACS-sorted Col-0 and Ler-0 pollen nuclei revealed decreased mCHH levels in Ler-0 VN, compared to Col-0 VN. **b**, Differentially methylated regions (DMRs) between Col-0 and Ler-0 VN were detected for CG and CHH methylation. mCHH hypomethylated DMR in Ler-0 VN overlapped primarily with TE features, particularly the superfamilies LTR/Gypsy and DNA/MuDR. **c**, Hypomethylated mCHH DMRs in Ler-0 VN overlapped with hypomethylated loci in *nrpd1a* (Pol IV) and *cmt2* mutant VN (both in Col-0 background). **d**, Small RNA in Col-0 pollen matching hypomethylated mCHH DMRs in Ler-0 VN are depleted in wild type Ler-0 pollen and dependent on Pol IV (*nrpdla*) (21/22 and 24nt). **e**, Genome browser tracks of CG, CHG and CHH methylation levels in the VN of Col-0, Ler-0, *nrpdla* and *cmt2* pollen, illustrating CMT2 and RdDM-targeted TEs where CHH methylation was lost in Ler-0 VN.

In Arabidopsis, most 20 to 22-nt miRNAs are processed by DICER-LIKE1 (DCL1) and loaded into ARGONAUTE1 (AGO1) in order to mediate post-transcriptional gene silencing (PTGS). We therefore crossed the GFP sensor with the null *dcl1-5* mutant allele and the strong hypomorphic *ago1-9* allele, and observed that GFP expression was restored in mutant pollen grains isolated from heterozygous plants, thus confirming that miR845 function in pollen requires the canonical miRNA pathway (Fig. 3a and Supplementary Fig. 1c). In somatic tissues, miRNA-directed biogenesis of secondary siRNAs from target transcripts requires either a “double hit” with two 21-nt miRNAs or a single hit with a 22-nt miRNA. Cleaved transcripts are converted into double stranded RNA (dsRNA) by the RNA-dependent RNA polymerase RDR6, and processed into 21- and 22-nt siRNAs by DCL4 and DCL2, respectively^25^. We used the miR845b-GFP sensor in *dcl1-5/+* and *dcl1/+,dcl2/dcl4* mutant backgrounds and purified wild-type, *dcl1*, *dcl2/4* and *dcl1/2/4* pollen by fluorescence-activated cell sorting (FACS) (Supplementary Fig. 1d). Small RNA sequencing of sorted pollen confirmed that most miRNAs were depleted in segregating *dcl1* mutant pollen, including miR845a and miR845b (Fig. 3c,d and Supplementary Fig. 1e), as well as abundant secondary siRNAs derived from the GFP transcript that were present in wild-type pollen but not in *dcl1*, *dcl24* and *dcl124* (Fig. 3b). However, loss of easiRNAs was observed only in *dcl2/4* pollen, but not in *dcl1* mutants (Fig. 3c), including 21/22-nt small RNAs produced from 5’ LTRs upstream of the PBS (miR845b target site) (Supplementary Fig. 4a). In somatic tissues, small RNAs matching to these LTRs are 24-nt in length and produced by Pol IV, RDR2, and DCL3^25^, raising the possibility that 21/22-nt easiRNA in pollen are dependent on Pol IV. Indeed, small RNA from *nrpd1a* mutant pollen lost siRNAs for the majority of TEs in all size classes (Supplementary Fig. 4b), indicating that easiRNA biogenesis in pollen at miR845 targets depends on Pol IV, DCL2 and DCL4. Similar pathways have been described under certain types of genotoxic stress, and may be relevant here^26,27^.

**Figure 3.**
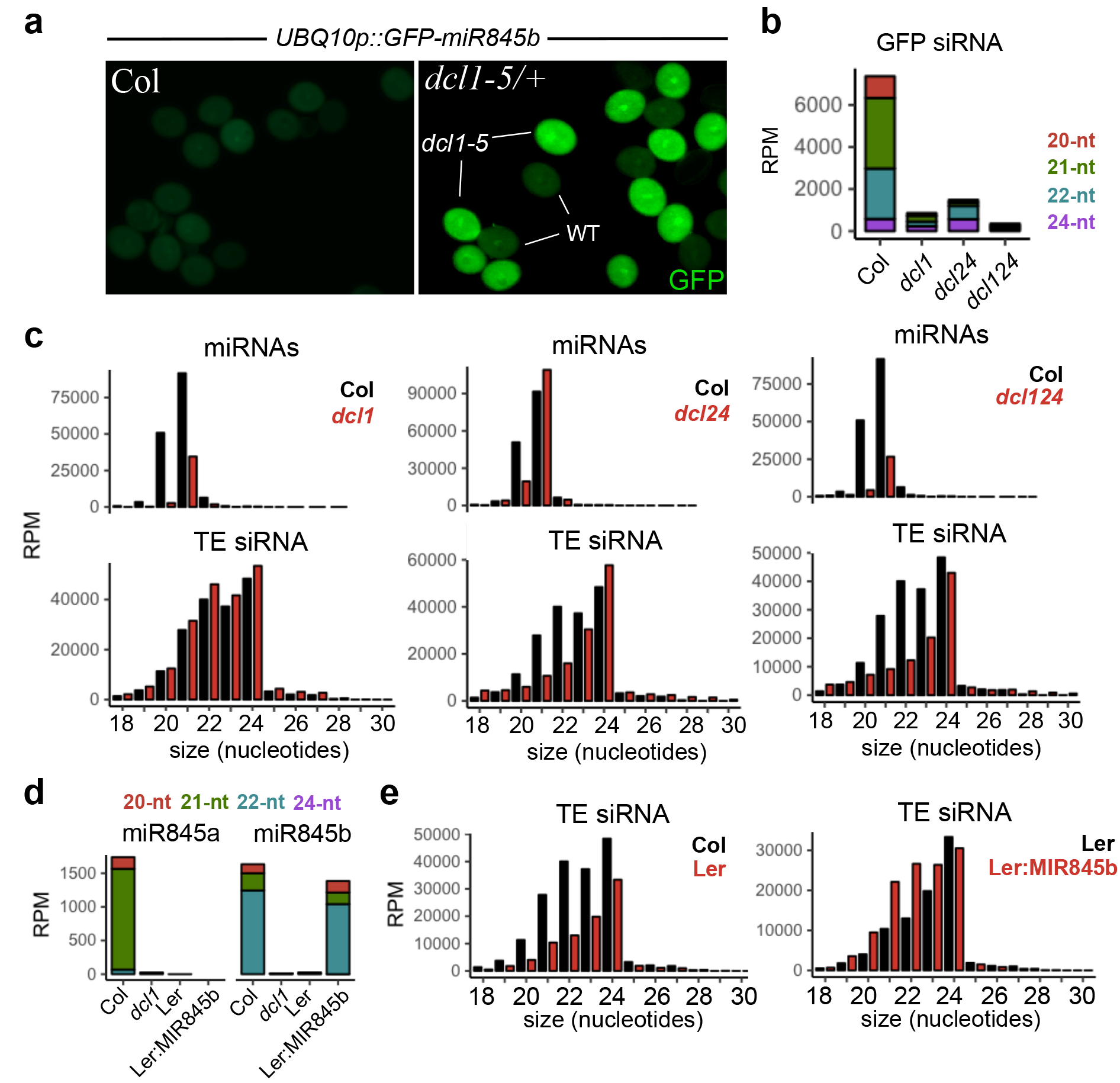
miR845b-dependent easiRNA biogenesis from transgenes and transposons. **a**, GFP sensor including 3’UTR with a miR845b target site and driven by the *UBIQUITIN10* (*UBQ10*) promoter in wild type and *dcl1-5/+* heterozygous background. GFP expression was restored in *dcl1* pollen, allowing FACS-purification of wild type, *dcl1, dcl2/4* and *dcl1/2/4* pollen grains. **b**, Loss of GFP siRNA was detected in *dcl1, dcl2/4* and *dcl1/2/4* pollen grains, indicating that miR845b triggers DCL2/4-dependent secondary siRNA from the GFP transgene. **c**, Small RNA sequencing from wild type and mutant FACS-sorted pollen revealed that 21- and 22-nt TE siRNA were lost in the *dcl2/4* mutants, while miRNAs were depleted in *dcl1*. **d**, miR845a and miR845b were depleted in *dcl1* mutant and Ler-0 pollen, but miR845bwas restored in transgenic Ler-0 plants expressing Col-MIR845b (Ler:MIR845b). **e**, 21- and 22-nt TE-derived siRNA levels were also depleted in wild-type Ler-0 pollen, but restored in transgenic Ler:MIR845b.

As these Pol IV-easiRNAs (21/22-nt) are not significantly depleted in *dcl1* pollen (Fig. 3c) where both miR845a and miR845b are down-regulated (Supplementary Fig. 1e), we hypothesized that miR845 contributes to Pol IV-easiRNA biogenesis either during meiosis or early at the onset of gametogenesis. In order to test this possibility, we cloned the Col-*MIR845b* locus and transformed Ler-0 wild-type plants. Strikingly, we observed strong and specific up-regulation of 21- and 22-nt TE-derived siRNAs, resembling wild-type Col-0 pollen (Fig. 3e). Interestingly, restored easiRNA biogenesis in Ler:MIR845b pollen did not result in significant changes in CHH methylation (Supplementary Fig. 5b), supporting the idea that easiRNAs in pollen do not modulate RdDM activity in the VN, because they are actively transported to the sperm cells^9,14^.

Parental small RNA differences can build strong barriers to hybridization^2^, and we have previously speculated that they might play a role in interspecific and interploid hybridization barriers in flowering plants^3^. Spontaneous chromosome doubling (polyploidization) is common in plants and is a major pathway towards reproductive isolation and speciation^4,28^. This is because hybrid seeds collapse as a result of unbalanced expression of imprinted genes in the endosperm, a phenomenon known as the “triploid block”. The “endosperm balance number” hypothesis further postulates the existence of specific loci responsible for seed collapse in different strains^4^. The triploid block can be conveniently demonstrated in Arabidopsis using the *omission of second division* (*osd1*) mutant that forms unreduced diploid male and female gametes that are self-fertile^29,30^. Diploid *osd1* pollen crossed to wild-type seed parents leads to the production of triploid seeds with tetraploid endosperm, that abort at high frequency depending on the genetic background^29,30^. In order to test whether miR845b-directed easiRNA biogenesis is involved in the triploid block response, we used a line carrying a T-DNA insertion at the *MIR845b* locus (*mir845b-1*) in Col-0 in which miR845b was down-regulated by roughly one half (Fig. 4a), and 21/22-nt easiRNAs were much lower than 24-nt siRNA (Fig. 4b), resembling *dcl2/4* mutants and wild type Ler-0 in this respect. Thus miR845b-directed easiRNA biogenesis is dose-sensitive, which is consistent with a role in endosperm balance. We next generated double mutants between *mir845b-1* and *osd1-1* in Col-0 to test whether loss of easiRNA in pollen has an effect on the triploid block. Pollinations of wild-type Col-0 seed parents with *osd1-1* homozygous pollen gives rise to approximately 5% of viable seeds, while pollinations with *osd1-1/mir845b-1* pollen had significantly increased seed viability at 35% (Fig. 4c). Interestingly, previous studies have shown that the triploid block response is much weaker in certain Arabidopsis accessions such as Ler-0^31,32^, where miR845-dependent easiRNAs are lost. Therefore, we crossed pollen from *osd1-2* (Ler-0 background) and *osd1-2* expressing Col-miR845b *(osd1-2:MIR845b)* to Ler-0 wild-type female parents. However, easiRNA up-regulation in Ler-0 diploid pollen (Fig. 4a,b) was not sufficient to restore the triploid block, since the levels of viable triploid seeds remained comparable (Fig. 4c). We conclude that miR845b-dependent easiRNAs are required but not sufficient for the triploid block, suggesting that there are additional regulatory mechanisms to balance parental gene dosage in Ler-0 endosperm, as previously reported^30,32^.

**Figure 4.**
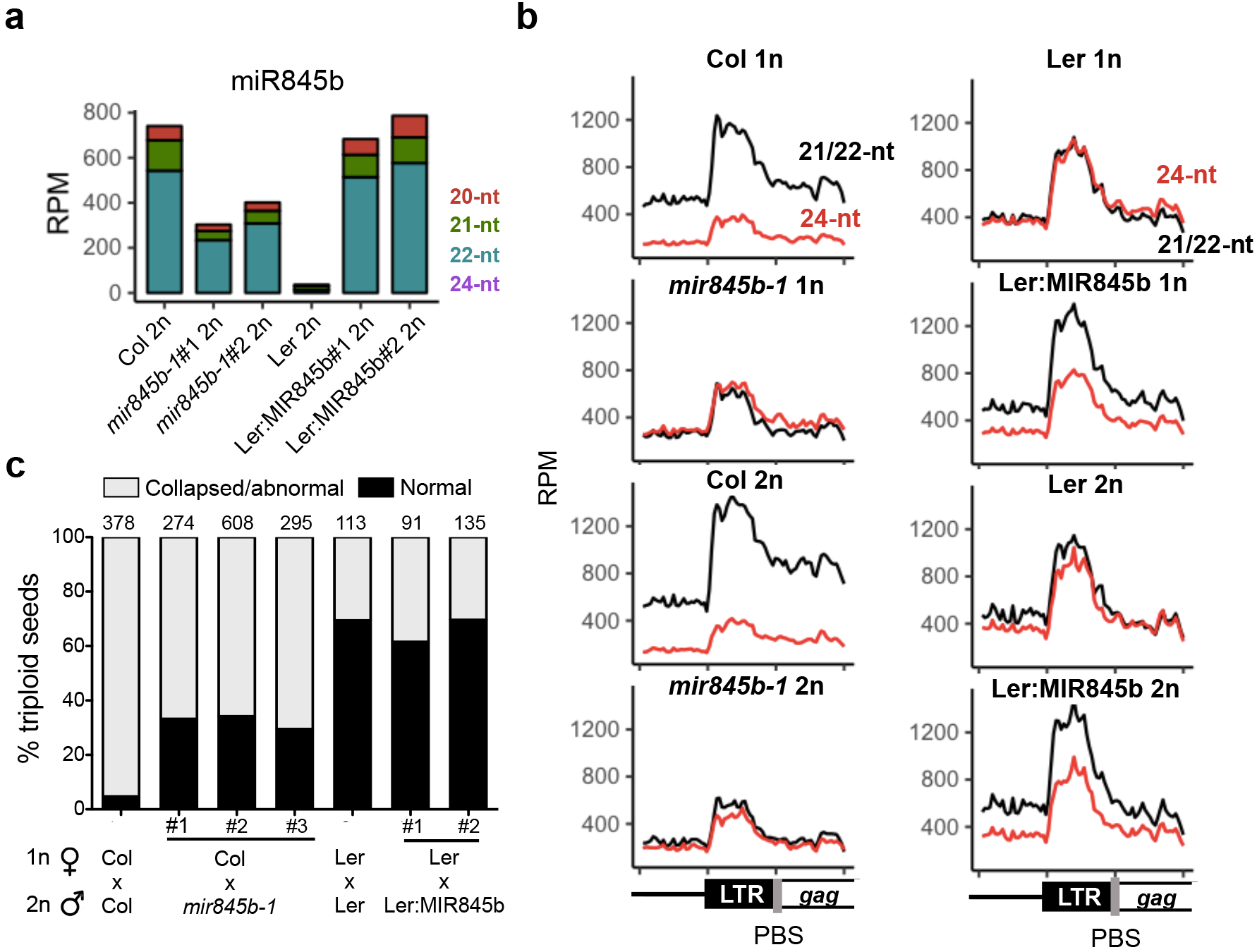
miR845b-dependent easiRNA is required for the triploid block. **a**, miR845b was down-regulated 2-fold in biological replicates of diploid pollen (2n) from a T-DNA insertion mutant at the *MIR845b* locus (*mir845b-1*), and abolished entirely in Ler-0. Expression of Col-*MIR845b* in Ler-0 2n pollen (*osdl-2* background, Ler:MIR845b) restored expression of miR845b in independent transgenic lines. **b**, TE-derived siRNAs were mapped to aligned 5’ regions (metaplots) of LTR *Gypsy* elements. 21/22-nt easiRNA were depleted in *mir845b-1* pollen and wild-type Ler-0 pollen compared to Col-0, but restored in Ler:MIR845b. These effects were even more pronounced in *osd1-2* diploid pollen (2n). **c**, Wild type Col-0 seed parents were pollinated with *osd1-1* diploid (2n) pollen leading to the production of triploid seeds that abort at high frequency. When *osd1-1/mir845b-1* double mutant (2n) pollen was used, seed viability increased to 35%. When *osd1-2* Ler-0 (2n) pollen was used, the triploid block was also partially suppressed with 30% abnormal or collapsed seeds. However, expression of Col-*MIR845b* in *osd1-2* Ler-0 2n pollen (Ler:MIR845b) was not sufficient to restore the triploid block.

In summary, paternally expressed miR845b stimulates dose-dependent biogenesis of 21/22-nt secondary siRNAs via Pol IV transcription, to mediate the triploid block response. These observations are consistent with previous results showing that the ratio of 21/22-nt vs 24-nt siRNA is much higher in interploid hybrid seeds with paternal excess^31–33^. Mechanistically, 21/22-nt easiRNAs might stabilize unbalanced genomic imprinting in interploid seeds, by promoting dose-dependent expression of imprinted genes in tetraploid endosperm. In previous studies, reduced levels of maternal 24-nt siRNAs resulted in up-regulation of maternally expressed genes (MEGs) but did not suppress triploid seed abortion^33^, while loss-of-function mutations in several paternally expressed genes (PEGs) are strong suppressors of the triploid block response^30,34^. Disrupted expression of PEGs and TEs is also associated with post-zygotic barriers in crosses between Arabidopsis species^35,36^, thus implying that up-regulation of PEGs is at least in part responsible for seed abortion upon interspecific and interploidy hybridizations.

One possibility is that paternal easiRNAs are involved in silencing MEGs, which include important components of the Polycomb Repressive Complex (PRC2) such as *MEDEA* (*MEA*) and *FERTILIZATION INDEPENDENT SEED 2* (*FIS2*). PEGs are silenced by PRC2-mediated H3K27me3 in the endosperm^37^, and so would be upregulated in triploid seeds. This idea is supported by the fact that MEGs accumulate 21/22-nt easiRNAs in interploid seeds with paternal excess, thus suggesting they are paternal in origin^33^ (Supplementary Fig. 6a-c), while maternal expression of *MEA* and *FIS2* is specifically repressed in these crosses^38^. Another possibility is that easiRNA promotes the expression of PEGs, by antagonizing RdDM activity at imprinted loci. In an accompanying paper, Martinez et al. show that paternal Pol IV mutations suppress the triploid block^39^, which is consistent with these mechanisms^15^. Our work reveals a key function for germline easiRNA in plants, suggesting parallels with germline piRNA in Drosophila that also control hybrid viability^2^, and 3’CCA tRF that regulate endogenous retroviruses in early mammalian development^20^. Dose-dependent regulation by miRNA might also contribute to “endosperm balance number”, which has been classically implicated in the triploid block^4^.

## Methods

### Plant material and growth conditions

Plants were grown under long day conditions at 22 °C. Seeds were always surface sterilized with sodium hypochlorite, sowed on Murashige and Skoog (MS) medium and stratified for 3 days at 4°C. Seedlings were transplanted to soil two weeks after germination and grown under long day conditions at 22 °C. We used the following Arabidopsis mutants: *dcl1-5* (CS16069), *ago1-9* ^40^ (originally in Ler, but introgressed in Col-0 for 6 generations of backcrossing), *mir845b-1* (SAIL_172_A08), *osd1-2* (GT21481), *nrpd1a-3* (Salk_128428) and *osd1-3* (Koncz collection) ^29^. The *mir845b-1* tetraploid plant was obtained by colchicine treatment as previously described ^41^. Additionally to Col-0 and Ler-0, we used the following ecotypes: Tsu-0 (CS1564), Kro-0 (CS1301), Bay-0 (CS954), Cvi-0 (CS902), RLD-1 (CS28687), Wei-0 (CS76301), Van-0 (CS28796), Sha (CS28737), Mt-0 (CS28502).

### Transgene cloning

The *UBQ10* promoter and miR845b target site promoter were cloned into a GFP reporter vector obtained from the VIB Department of Plant Systems Biology, UGent, by using Gateway technology (Life Technologies). Ectopic expression of MIR845a and MIR845b in Col-0 was performed by gateway cloning both MIR genes into pB2GW7. Expression of Col-MIR845b in Ler-0 was performed by PCR amplification of 1kb fragment of genomic DNA flanking MIR845b in Col-0, which was cloned into pMDC123 by Gateway cloning system. Primers used in this study are listed in Supplementary Table 4.

### Pollen collection and FACS

Pollen was collected in 1.5mL eppendorf tubes by vortexing open flowers in pollen extraction buffer (PEB, 10 mM CaCl_2_, 2 mM MES, 1 mM KCl, 1% H_3_BO_3_, 10%) for 3 min ^42^, followed by filtration through a 30um mesh (Partec/Sysmex) and centrifugation at 5,000g for 1 min. Pollen was suspended in 50ul of PEB, immediately frozen in liquid nitrogen and stored at - 80°C until RNA extraction. Purification of pollen expressing the GFP-miR845b sensor in *dcl1-5/+* and *dcl1/+,dcl2/dcl4* mutants was performed by vortexing open flowers in 1 mL of PEB and filtering through a 30um mesh before FACS. Pollen population was identified in SSC/FSC scatter plots, and GFP fluorescence was analyzed by excitation with a 488nm laser and detected with a 530/30 bandpass filter (Supplementary Fig. 1d).

### RNA purification, qPCR analysis and sequencing

Pollen total RNA was extracted using Trizol LS reagent (Life Technologies) by vortexing with glass beads for 5 minutes, and concentrated with Direct-zol columns (Zymo Research). miRNA-qPCR was performed using the Quantimir kit (System Biosciences) following manufacturer instructions. Libraries for RNA sequencing from Col-0 and Ler-0 pollen were prepared using rRNA-depleted total RNA samples and the ScriptSeq-v2 RNA-Seq Library Prep kit (Epicenter/Illumina), following manufacturer instructions. cDNA libraries were sequenced on a HiSeq2500 instrument. FastQ files were processed and mapped to TAIR10 genome using SubRead ^43^, normalized by calculating the number of paired-end reads per kilobase of transcript per million of mapped reads (FPKM), and analyzed using R scripts. Small RNAs were purified by running total RNA on acrylamide gels (15% polyacrylamide, 7M urea) and performing size selection of approximately 18 to 30-nt region using a small RNA ladder (Zymo Research). The small RNA fraction was isolated from acrylamide gel plugs by grinding with a plastic pestle in Trizol LS (Life Technologies), and concentrated using Direct-zol columns (Zymo Research). Libraries were prepared using the TruSeq Small RNA Sample Preparation Kit (Illumina) following manufacturer instructions, barcoded and sequenced in Illumina HiSeq 2500, NextSeq500 or MiSeq platforms depending on sample pooling strategies and desired sequencing depth. After de-multiplexing, 36-nt reads were pre-processed by trimming the 3’ adapter and filtering collapsed reads according to length and quality. Filtered reads were mapped to the Arabidopsis TAIR10 genome annotation (or GFP transgene) with bowtie reporting all multi-mappers. Only perfect match reads were used for down-stream analysis, and reads mapped to multiple genomic locations where normalized by dividing non-redundant read counts by the number of genomic hits, and subsequently calculating the number of reads per million of filtered (18-30nt) and perfectly mapped reads. Additional downstream analyses were performed using R scripts. A summary of all small RNA sequencing data generated in this study is presented in Supplementary Table 3. Inflorescence small RNA data was previously reported^21^.

### Bisulfite sequencing and DNA methylation analysis

Pollen nuclei were isolated as previously described^42,44^. Approximately 50.000 nuclei were purified by FACS and used to construct sequencing libraries of bisulfite-treated DNA using the Pico Methyl-Seq kit (Zymo Research), according to manufacturer instructions. Non-directional libraries were sequenced on HiSeq and NextSeq platforms. Single end 100 and/or 76 reads were preprocessed using Trimmomatic ^45^ to remove adapters, trim the first 10 nucleotides and split the read in half. Preprocessed C/T- and G/A-converted reads were mapped to C/T- and G/A-converted TAIR10 genome allowing two mismatches. Perl and Python scripts were used to recover the sequence of each mapped read and calculate methylation at each individual cytosine covered by 3 or more reads. Differentially methylated regions (DMRs) were defined as 100bp bins containing at least 4, 6 or 8 differentially methylated mCGs, mCHGs or mCHHs, respectively, and absolute methylation difference of 0.35. DMRs localizing 200bp of each other were merged. A summary of all genome-wide bisulfite sequencing data generated in this study is presented in table S4. Bisulfite sequencing data of *cmt2* mutant VN was previously reported ^23^.

## Acknowledgements

We thank all members of the Martienssen laboratory for discussions. Research in the Martienssen laboratory is supported by the US National Institutes of Health (NIH) grant R01 GM067014, The National Science Foundation Plant Genome Research Program, and by the Howard Hughes Medical Institute and Gordon and Betty Moore Foundation. The authors acknowledge assistance from the Cold Spring Harbor Laboratory Shared Resources, which are funded in part by the Cancer Center Support Grant (5PP30CA045508). Research in the Köhler laboratory was supported by a European Research Council Starting Independent Researcher grant, a grant from the Swedish Science Foundation and a grant from the Knut and Alice Wallenberg Foundation (to C.K.). G.M. was supported by a Marie Curie IOF Postdoctoral Fellowship (PIOF-GA-2012-330069). Sequencing for the Köhler laboratory was performed by the SNP&SEQ Technology Platform, Science for Life Laboratory at Uppsala University, a national infrastructure supported by the Swedish Research Council (VRRFI) and the Knut and Alice Wallenberg Foundation.

## Author contributions

F.B., C.K. and R.A.M designed the study. F.B., J.S.P., F.V.E., P.W. and G.M performed experiments, and F.B. analyzed the data. All authors contributed with ideas and discussion. F.B. and R.A.M prepared the manuscript.

## Supplementary information

**Supplementary Figure 1.**
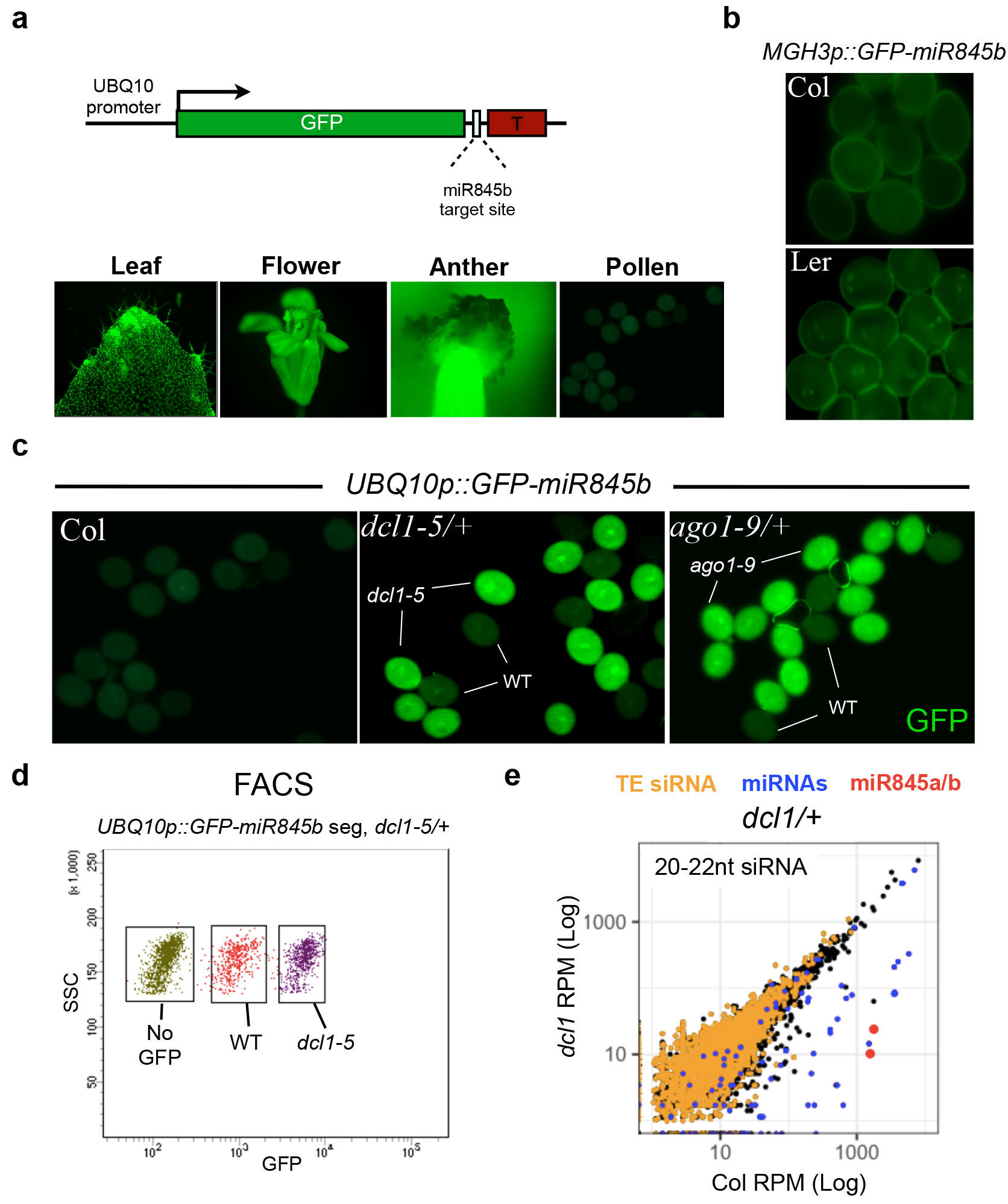
miR845b activity in somatic tissues and pollen. **a**, Expression of a GFP reporter with a miR845b target site in the 3’UTR was driven by the *UBIQUITIN10* (*UBQ10*) promoter and visualized in wild-type somatic tissues. GFP silencing was observed only in pollen. **b**, miR845b was active in Col-0 sperm cells, as expression of the GFP-miR845b construct driven the sperm-specific MGH3 promoter was silenced. GFP expression in sperm cells was observed in Ler-0 pollen, since miR845b is not expressed in this ecotype. c, GFP sensor was silenced in wild-type pollen, but not in *ago1-9* and *dcl1-5* pollen. **d**, The GFP sensor in *dcl1-5/+* heterozygotes was used to sort wild-type (GFP negative) and *dcl1-5* (GFP positive) pollen. **e**, Small RNA sequencing of FACS-sorted pollen populations revealed that most miRNAs, including miR845a and miR845b, were depleted in *dcl1-5* pollen.

**Supplementary Figure 2.**
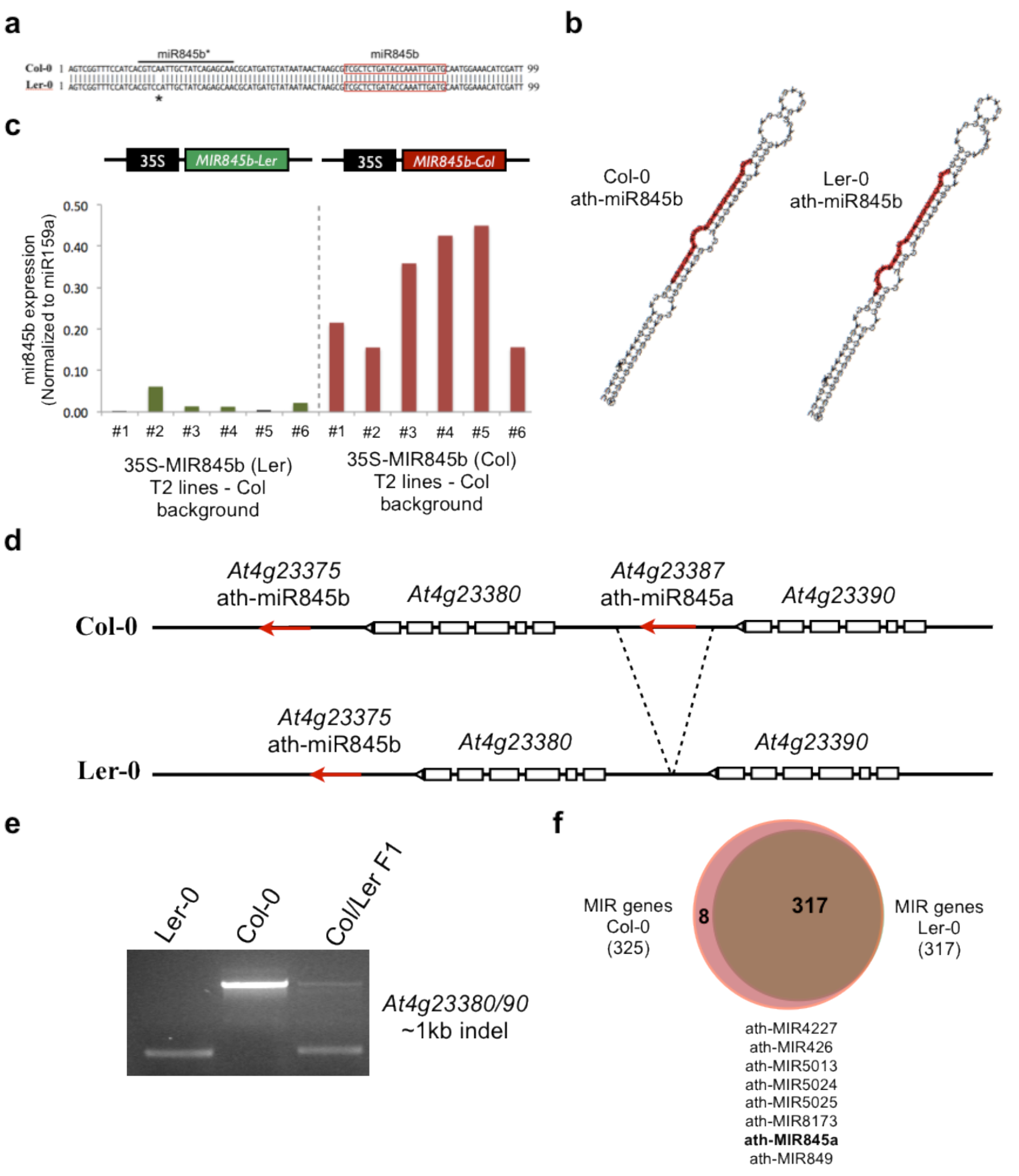
Natural variation in miR845 biogenesis in Arabidopsis Col-0 and Ler-0. **a**, A Single Nucleotide Polymorphism (SNP) in the miR845b* region of MIR845b between Col-0 and Ler-0. **b**, SNPs in MIR845b change the predicted structure of the MIR845b stem-loop, which could impair processing by DCL1. **c**, Ectopic expression of Col-MIR845b and Ler-MIR845b in leaves driven by the 35S promoter confirms that the Ler-MIR845b is not efficiently processed in several independent transgenic lines. **d**, *MIR845a* gene was identified in Col-0 (TAIR10 annotation). In Ler-0, there is a 1kb deletion at the *MIR845a* locus^46^. **e**, This indel was confirmed by PCR in Col-0, Ler-0 and Col/Ler-0 F1 hybrid. **f**, Venn diagram shows that in addition to *MIR845a*, there are 7 additional MIRNA genes annotated in Col-0 that are not present in Ler-0.

**Supplementary Figure 3.**
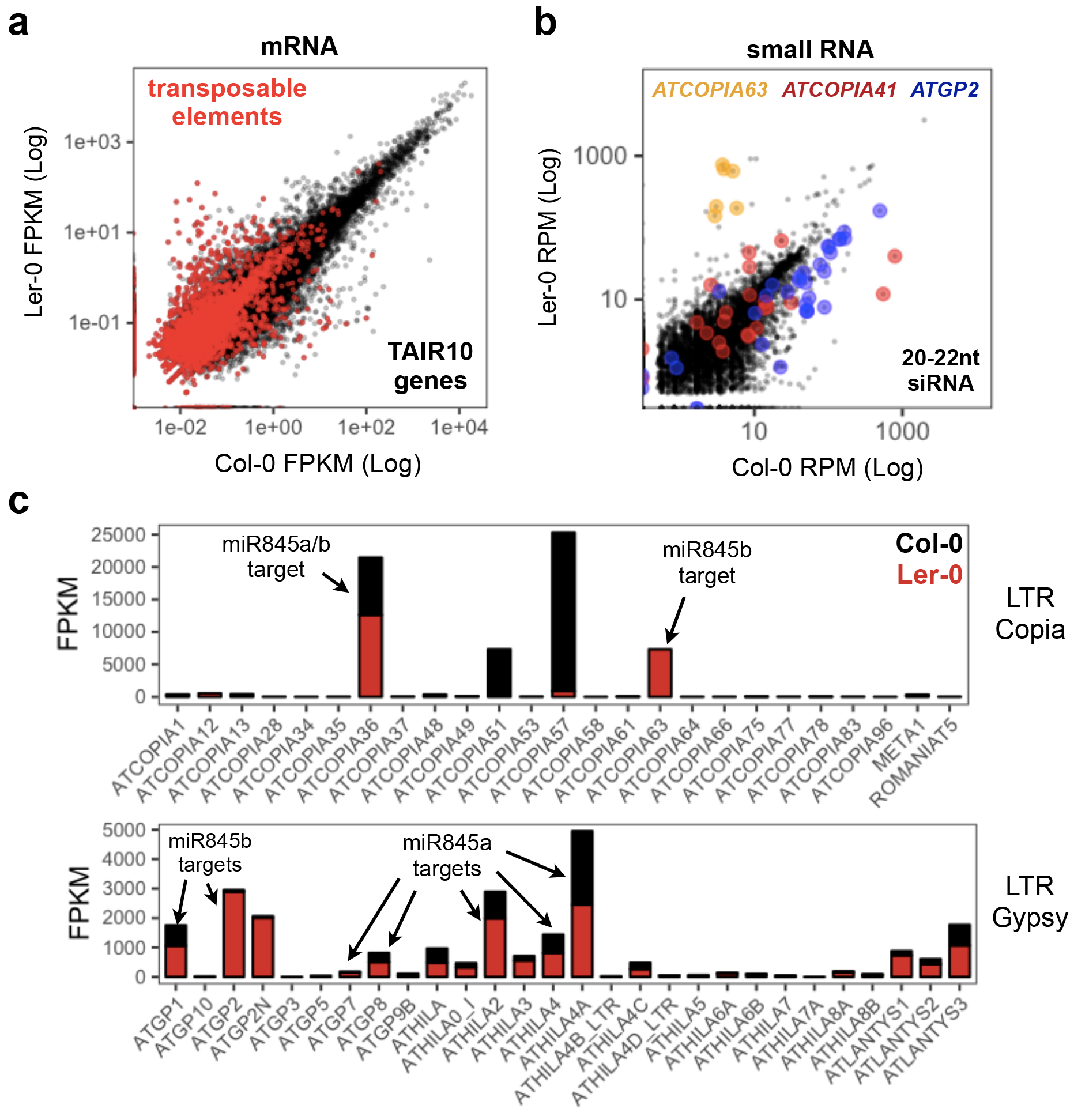
Natural variation in transposon expression and easiRNA biogenesis in Col-0 and Ler-0 pollen. **a**, Col-0 and Ler-0 pollen transcriptomes revealed that TE transcripts were overall more abundant in Ler-0 pollen. **b**, These differences are reflected in the accumulation of easiRNAs in pollen isolated from each ecotype. For example, miR845-targeted TEs *ATGP2* and *ATCOPIA41* accumulate easiRNA in Col pollen, while *ATCOPIA63* is only expressed in Ler-0 where it accumulates easiRNA, presumably via alternative miRNA. **c**, LTR retrotransposon (*Gypsy* and *Copia*) that are predicted miR845 targets show enriched expression in Ler-0 pollen.

**Supplementary Figure 4.**
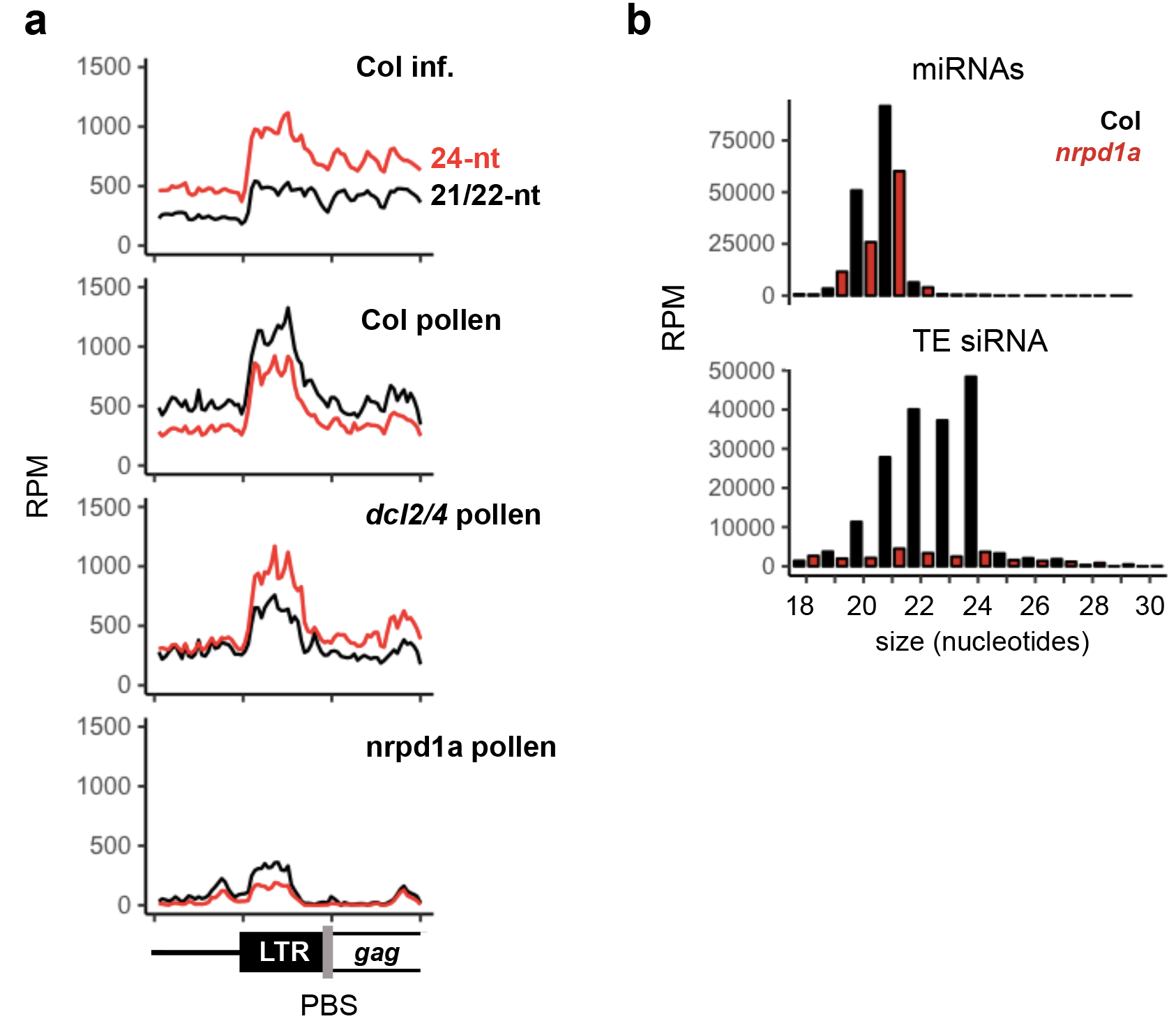
miR845b-dependent easiRNAs at the LTRs of *Gypsy* retrotransposons. **a**, Normalized small RNA reads were mapped to aligned 5’ regions (metaplots) of all annotated *Gypsy* elements, including 2kb upstream and 4kb downstream of the LTR. This includes the 5’ LTR (~2kb), PBS (miR845b target site) and part of the *gag* gene. 21/22-nt siRNAs were more abundant in wild-type pollen but depleted in the *dcl2/4* mutant pollen. Both 21/22- and 24-nt siRNAs were depleted *nrpd1a* (Pol IV) mutant pollen. **b**, Most small RNAs matching to TEs were lost in the *nrpd1a* mutant pollen. Many miRNA were also reduced by 2-fold, likely due to destabilization of ARGONAUTE proteins in *nrpd1a* mutant pollen.

**Supplementary Figure 5.**
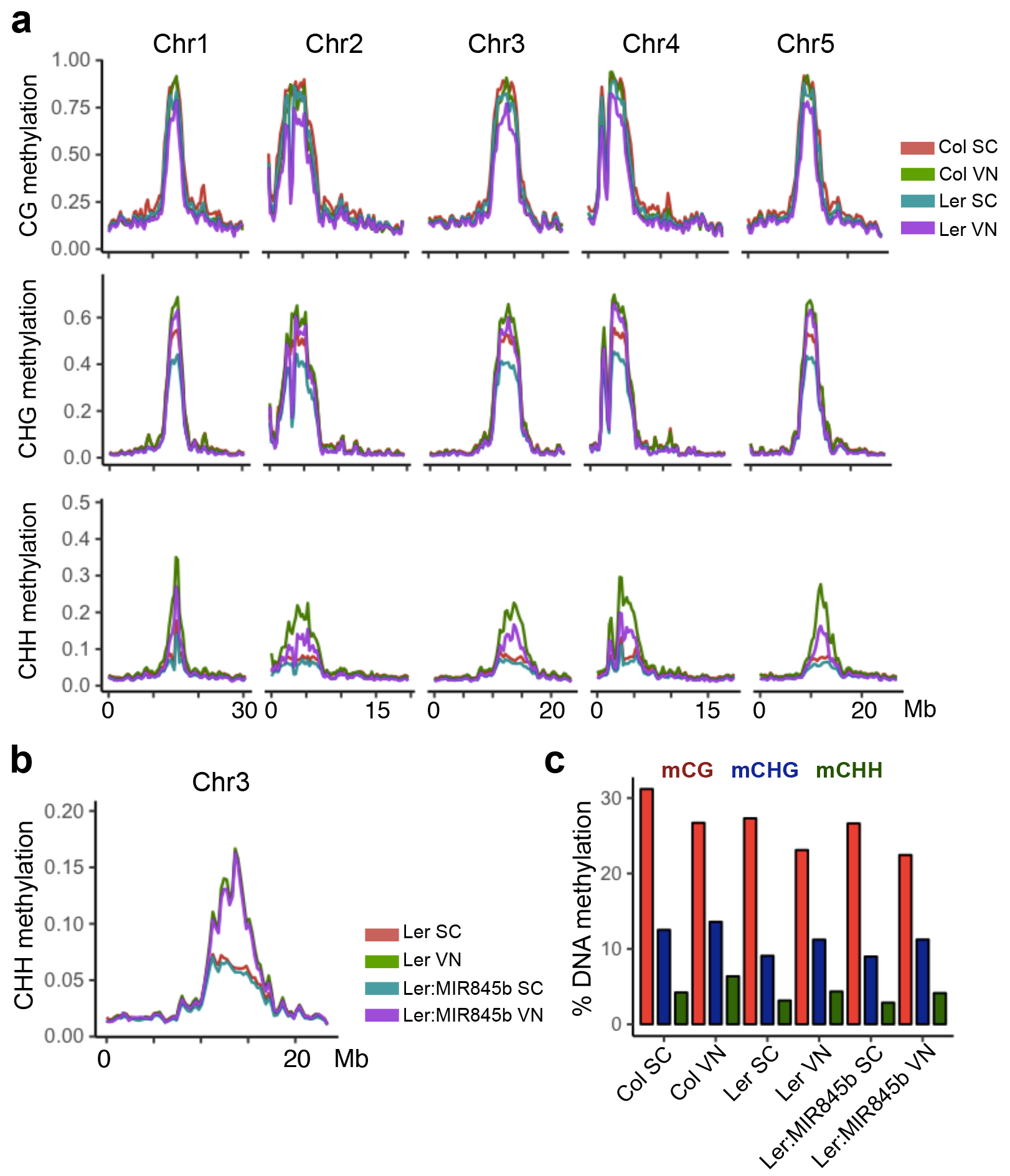
DNA methylation levels in Col and Ler pollen nuclei. **a**, Average DNA methylation levels in Col-0 and Ler-0 sperm (SC) and vegetative nuclei (VN) were plotted in 100kb windows on all 5 chromosomes in Arabidopsis. **b**, DNA methylation in the CHH context in SC and VN was not significantly changed in Ler-0 transgenics expressing Col-*MIR845b* (Ler:MIR845b) in pollen. **c**, Histogram representation of average DNA methylation percentages in Col, Ler-0 and Ler:MIR845b SC and VN nuclei.

**Supplementary Figure 6.**
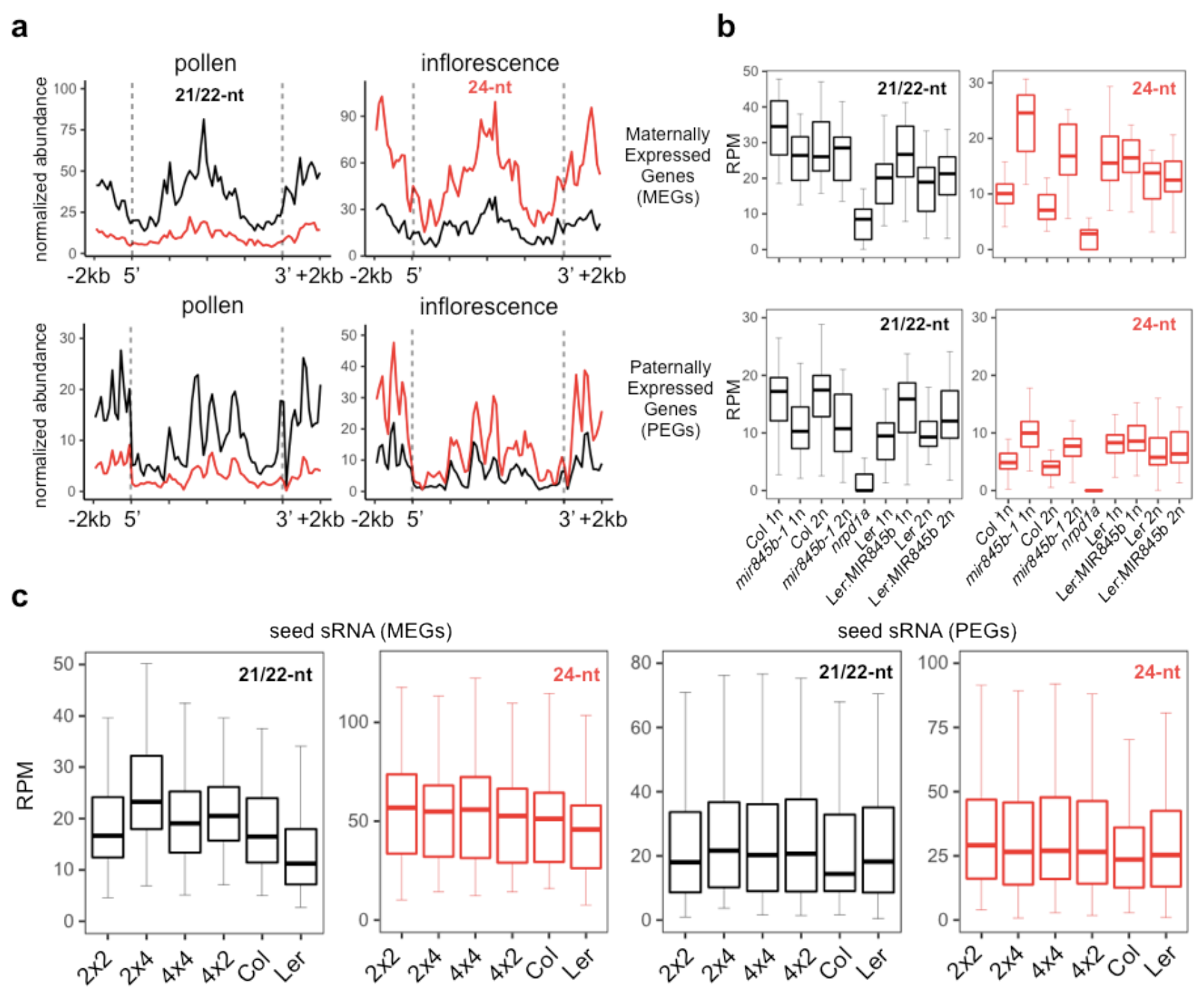
Small RNA abundances at MEGs and PEGs. **a**, Small RNA reads were mapped to aligned metaplots of maternally (MEGs) and paternally expressed genes (PEGs)^22^, including 2kb upstream and downstream of annotated coding regions, and normalized by total mapped reads. MEGs and PEGs accumulate 21/22nt easiRNA in pollen. **b**, Boxplots of normalized siRNA abundances matching 2kb regions flanking MEGs and PEGs in haploid (1n) and diploid (2n) pollen isolated from wild type Col-0, *mir845b-1*, *nrpd1a-3*, wild type Ler-0 and Ler-0 transgenics expressing Col-miR845b (Ler:MIR845b). 21/22nt easiRNA matching MEGs and PEGs depend on miR845b and polIV. **c**, Boxplots of published small RNA datasets from diploid (2×2), tetraploid (4×4) and interploid (2×4) seeds ^33^ and from Col-0 and Ler-0 seeds ^22^. MEGs accumulated higher levels of 21/22-nt easiRNA in interploid (2×4) seed.

**Supplementary Table 1.**
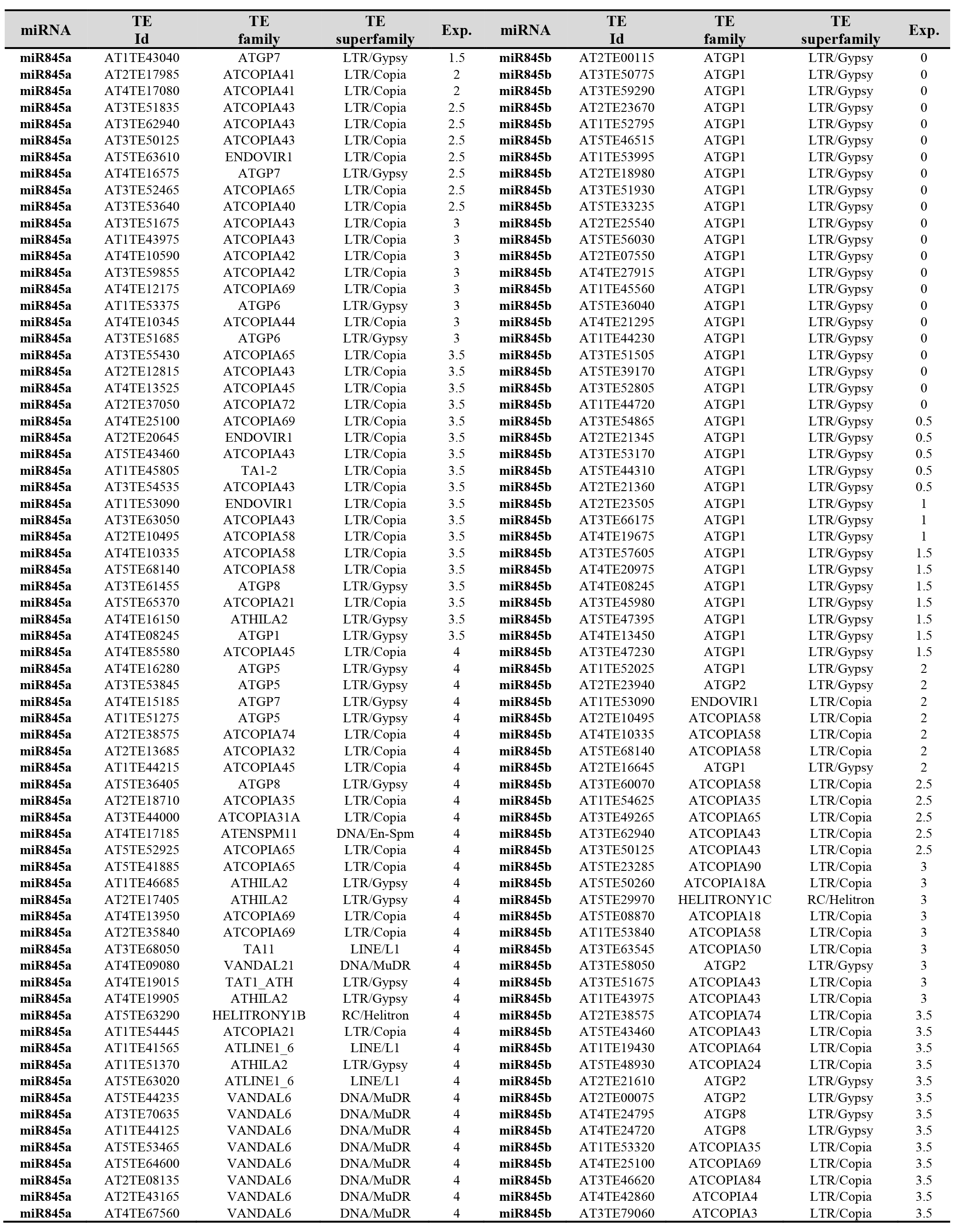
miR845 target sites in retrotransposons were predicted using the psRNATarget software (2011 release) (*49*), using default settings for all parameters, except the maximum expectation value (Exp < 5.0).

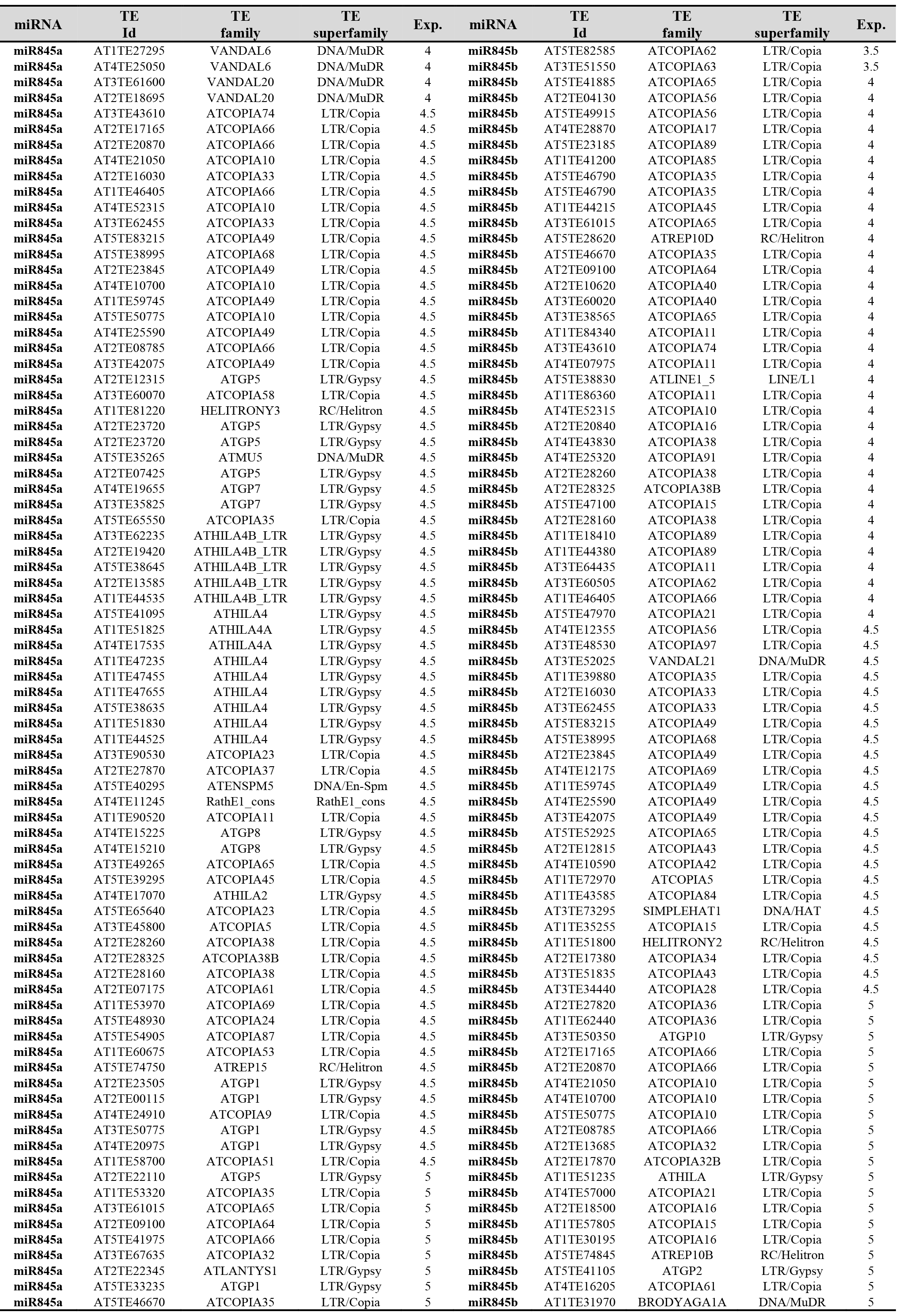

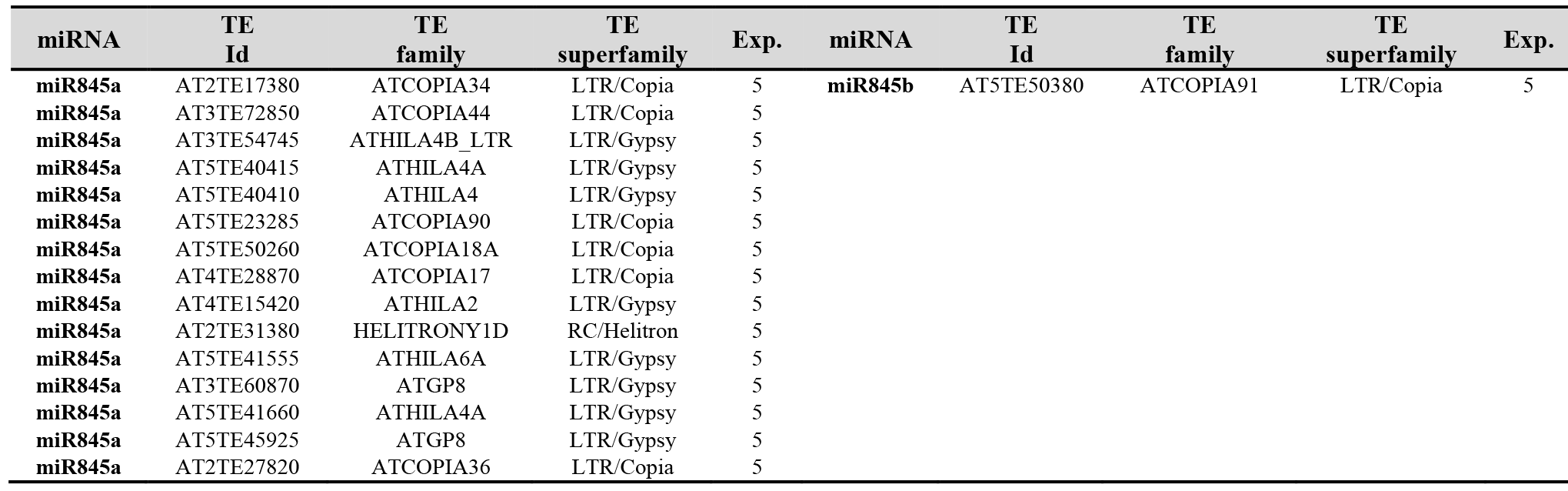

**Supplementary Table 2.**
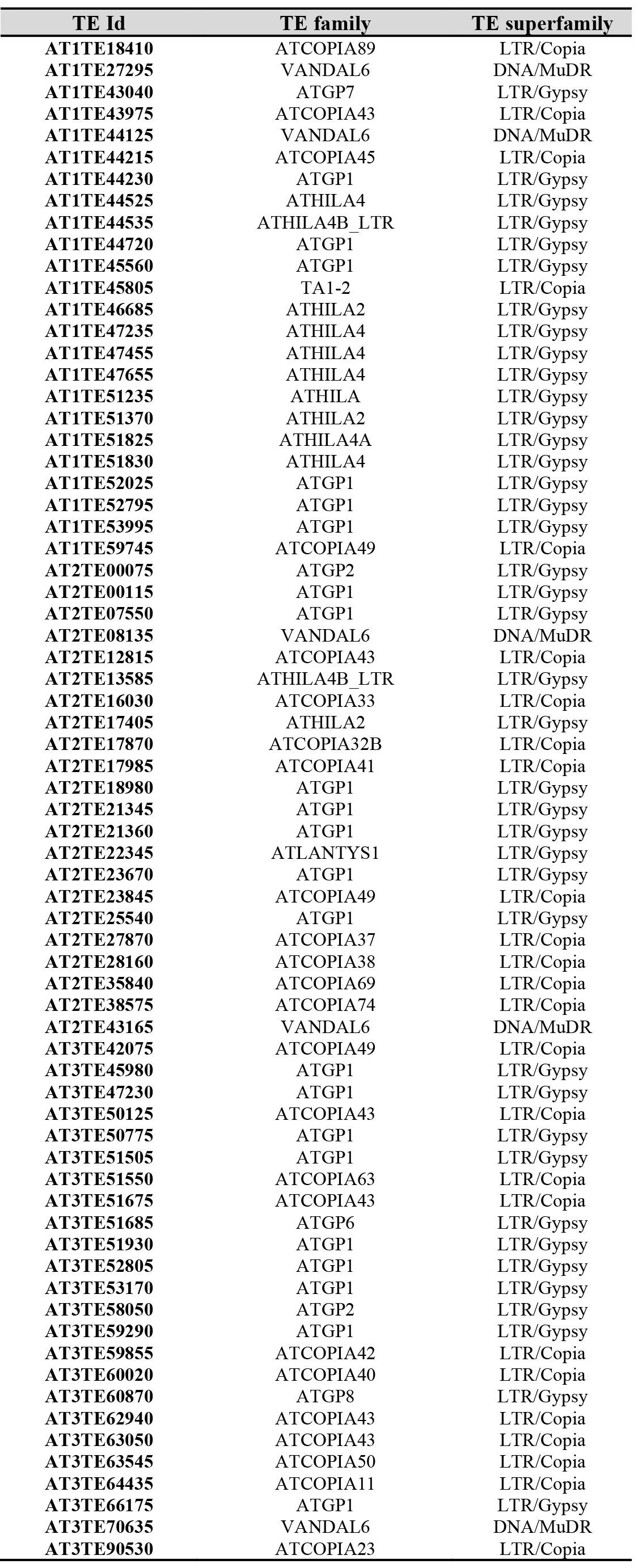
Predicted miR845 targets overlapping with mCHH hypomethylated DMRs in Ler-0 pollen.

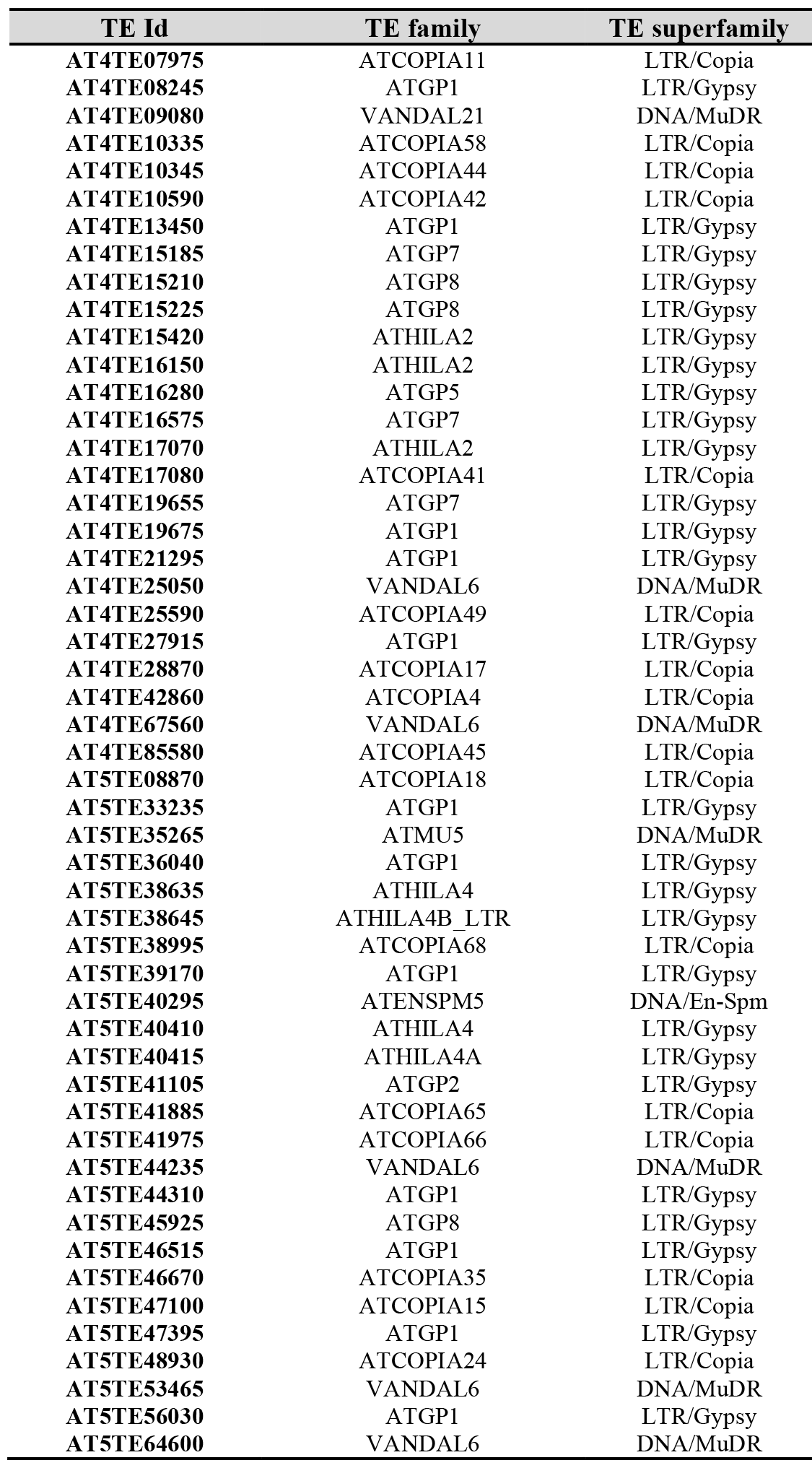

**Supplementary Table 3.** Summary of small RNA and whole genome bisulfite sequencing. **a**, Trimmed reads (18-30 nucleotides) were mapped to TAIR10 genome using bowtie (-v 1). **b**, Mean genomic coverage and methylation percentages for CG, CHG and CHH context. Cytosines covered by 3 or more reads were used to calculate methylation percentages.

| a, Small RNA libraries |  |  |  |  |  |  |  |
| --- | --- | --- | --- | --- | --- | --- | --- |
| sample | ecotype | trimmed<br>(18-30nt)<br>total reads | trimmed<br>(18-30nt)<br>mapped reads | mapping efficiency<br>% |  |  |  |
| Col pollen FACS | Col | 2,960,479 | 2,886,915 | 97.5 |  |  |  |
| *dcl1* pollen FACS | Col | 4,547,667 | 4,452,340 | 97.9 |  |  |  |
| *dc2/4* pollen FACS | Col | 1,728,590 | 1,622,842 | 93.9 |  |  |  |
| *dcl1/2/4* pollen FACS | Col | 1,338,696 | 1,241,077 | 92.7 |  |  |  |
| Ler pollen | Ler | 17,107,192 | 15,452,395 | 90.3 |  |  |  |
| Ler:MIR845b pollen | Ler | 20,778,823 | 19,321,088 | 93.0 |  |  |  |
| *osd1-2* pollen | Ler | 12,884,293 | 11,743,763 | 91.1 |  |  |  |
| *osd1-2*:MIR845b pollen | Ler | 19,099,994 | 18,190,694 | 95.2 |  |  |  |
| *nrpd1a-3* pollen | Col | 769,276 | 705,167 | 91.7 |  |  |  |
| *mir845b-1* 1n rep 1 pollen | Col | 10,345,592 | 10,016,713 | 96.8 |  |  |  |
| *mir845b-1* 1n rep 2 pollen | Col | 13,461,019 | 13,042,057 | 96.9 |  |  |  |
| *mir845b-1* 2n rep 1 pollen | Col | 6,989,445 | 6,328,951 | 90.6 |  |  |  |
| *mir845b-1* 2n rep 2 pollen | Col | 13,636,290 | 10,788,076 | 79.1 |  |  |  |
| Col leaf | Col | 2,866,399 | 2,811,539 | 98.1 |  |  |  |
| b, Bisulfite libraries |  |  |  |  |  |  |  |
| sample | ecotype | mean<br>coverage<br>CG | mean<br>coverage<br>CHG | mean<br>coverage<br>CHH | %CG<br>(>=3) | %CHG<br>(>=3) | %CHH<br>(>=3) |
| Col SC | Col | 14.4 | 13.4 | 11.0 | 31.6 | 12.2 | 4.3 |
| Col VN | Col | 15.1 | 14.1 | 11.8 | 27.5 | 13.1 | 6.3 |
| Ler SC | Ler | 14.5 | 14.4 | 13.2 | 26.2 | 8.1 | 3.1 |
| Ler VN | Ler | 9.9 | 9.1 | 8.2 | 22.2 | 10.0 | 4.3 |
| Ler:MIR845b SC | Ler | 9.6 | 9.1 | 8.4 | 25.4 | 8.0 | 2.8 |
| Ler:MIR845b VN | Ler | 10.9 | 11.0 | 9.7 | 21.5 | 9.9 | 4.0 |
| *nrpd1a-3* VN | Col | 6.0 | 5.6 | 4.7 | 28.1 | 13.7 | 5.9 |

**Supplementary Table 4.**
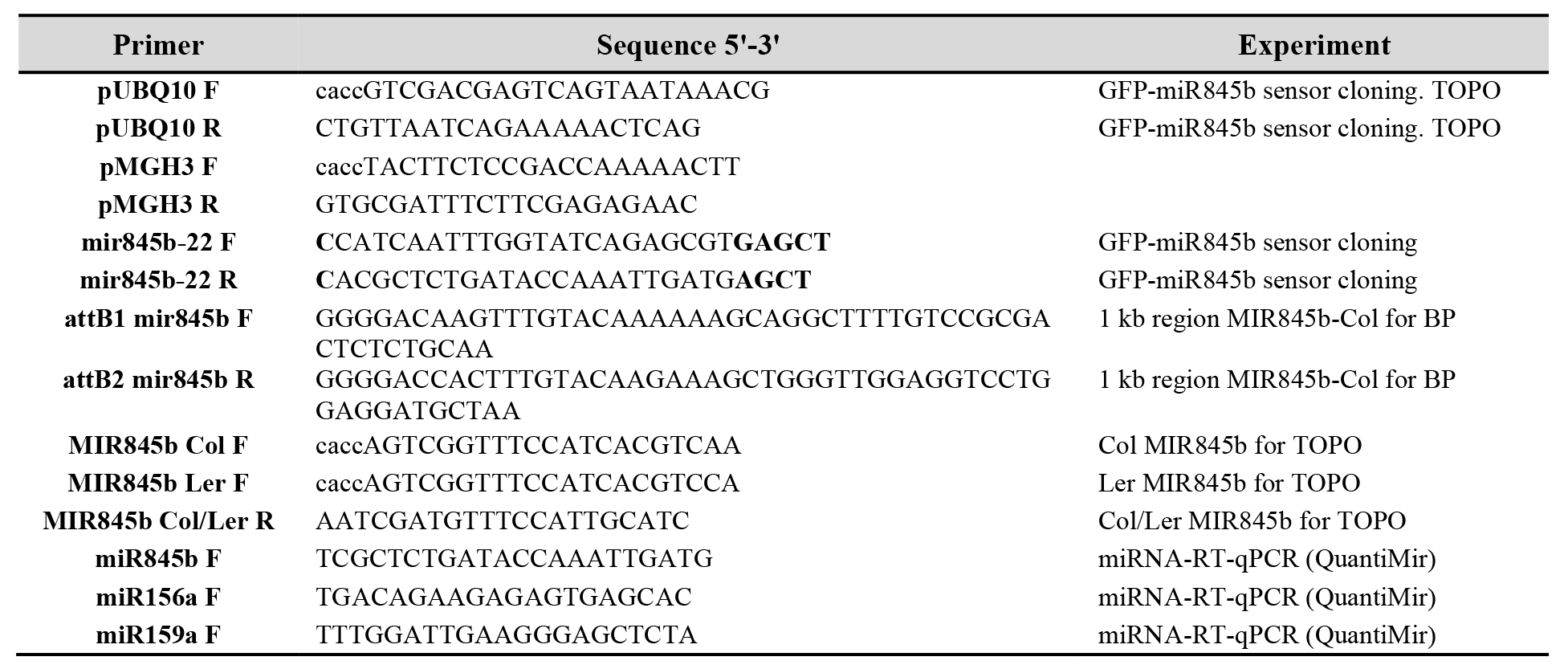
Oligonucleotide primers used in this study.

